# Natural selection can favor ratchet robustness over mutational robustness

**DOI:** 10.1101/121087

**Authors:** Yinghong Lan, Aaron Trout, Daniel Michael Weinreich, Christopher Scott Wylie

**Affiliations:** Department of Ecology and Evolutionary Biology, and Center for Computational Molecular Biology, Brown University, Providence, RI, USA; Department of Mathematics, Chatham University, Pittsburgh, PA, USA

**Keywords:** natural selection, ratchet robustness, mutational robustness, fitness landscape, Muller’s ratchet

## Abstract

The vast majority of fitness-affecting mutations are deleterious. How natural populations evolve to cope is a question of fundamental interest. Previous studies have reported the evolution of mutational robustness, that is, natural selection favoring populations with less deleterious mutations. By definition, mutational robustness provides a short-term fitness advantage. However, this overlooks the fact that mutational robustness decreases finite asexual populations’ ability to purge recurrent deleterious mutations. Thus, mutational robustness also results in higher risk of long-term extinction by Muller’s ratchet. Here, we explore the tension between short- and long- term response to deleterious mutations. We first show that populations can resist the ratchet if either the selection coefficient or the ratio of beneficial to deleterious mutations increases as fitness declines. We designate these properties as ratchet robustness, which fundamentally reflects a negative feedback between mutation rate and the tendency to accumulate more mutations. We also find in simulations that populations can evolve ratchet robustness when challenged by deleterious mutations. We conclude that mutational robustness cannot be selected for in the long term, but it can be favored in the short-term, purely because of temporary fitness advantage. We also discuss other potential causes of mutational robustness in nature.

## Introduction

The vast majority of fitness-affecting mutations are deleterious. How natural populations evolve to cope is a question of fundamental interest. One widespread hypothesis proposes the evolution of mutational robustness, which would mean that deleterious mutations evolve to become less deleterious(Krakauer and Plotkin 2001; Wilke and Adami 2003). Intuitively, it may seem that organisms with higher mutational robustness will have a fitness advantage over those with lower mutational robustness. Taking that intuition one step further, organisms with higher deleterious mutation rate, i.e., more frequent occurrences of deleterious mutations, might be expected to evolve greater mutational robustness. And indeed, there exists a substantial body of literature consistent with that idea (**Box 1**).

### Box 1: Literature on the evolution of mutational robustness

Two computational studies (van Nimwegen et al. 1999; Wilke et al. 2001) have addressed the possibility of evolution of mutational robustness. Using Avida, a platform for conducting in silico evolutionary experiments (Ofria et al. 2009), Wilke et al. (2001) evolved populations founded with the same ancestor under high and low mutation rates. Then they competed the two evolved lineages against each other under different mutation rates. They found that the lineage evolved under high mutation rate, albeit having lower fitness, wins under higher mutation rate. Their interpretation was that that the lineage adapted to high mutation rate has higher mutation robustness. Two experimental studies using a viroid (Codoner et al. 2006) or an RNA virus (Sanjuan et al. 2007) similarly found that in competition at high mutation rate, populations evolved at high mutation rate displaced those evolved at low mutation rate.

Working in the neutral network framework, van Nimwegen et al. (1999) explored a model in which mutations are either neutral or lethal. In this case, mutational robustness of each genotype is measured by its proportion of neutral mutations. They found that as mutation rate increases, populations will evolve to be composed of genotypes with fewer lethal mutations, i.e., with higher mutational robustness.

However, despite the intuitive appeal of the foregoing, it overlooks important components of population genetics theory. Firstly, in infinite populations, where genetic drift can be neglected, equilibrium mean fitness depends only on deleterious mutation rate (Haldane 1937; Kimura and Maruyama 1966). Thus, mutational robustness, i.e., the reduced effect of deleterious mutations, does not confer any long-term fitness advantage. Moreover, finite asexual populations risk Muller’s ratchet (Muller 1964; Haigh 1978), meaning that deleterious mutations accumulate at a rate so fast that selection is overwhelmed. In other words, besides the immediate fitness cost of deleterious mutations, Muller’s ratchet represents a longer-term challenge to evolving populations. A previous study (Goyal et al. 2012) provides the quantitative parametric boundary above which populations persist at Mutation-Selection-Drift Equilibrium (MSDE) (Wager and Gabriel 1990; Schultz and Lynch 1997; Poon and Otto 2000), but below which they succumb to the ratchet. In doing so, this study demonstrates that finite populations are less susceptible to Muller’s ratchet when mutation rate is lower, population size is greater, selection coefficient is higher, or the ratio of beneficial to deleterious mutation rates is larger.

A fundamentally important question then arises as to how populations evolve in the face of both short- and long-term challenges brought by deleterious mutations? Trivially, the evolution of lower mutation rate will alleviate both concerns. Therefore, throughout this paper we hold mutation rate as an extrinsic factor and observe other responses of evolving populations. Furthermore, we hold population size constant, i.e., we assume soft selection. This approach is conservative, because decreasing population size renders the population more prone to Muller’s ratchet, ramifications of which have already been studied in the context of mutational meltdown (Lynch et al. 1993; Bull et al. 2005). Consequently, we focus only on the influence of selection coefficient and the ratio of beneficial to deleterious mutation rates. As a complement to the phrase “mutational robustness”, we correspondingly designate increases in either of these properties as conferring “ratchet robustness.” (Recently Labar and Adami 2016 introduced the phrase drift robustness for a similar phenomenon. However, we prefer ratchet robustness, since it more precisely describes the underlying population genetic risk against which robustness is conferred.)

The fitness landscape provides a unifying framework for exploring both mutational robustness and ratchet robustness. First define genotype space, where spatially adjacent genotypes are mutationally adjacent, and then project the fitness of each genotype over the genotype space. Since mutational robustness means deleterious mutations have smaller selection coefficients, regions of the landscape with higher mutational robustness are “flatter”. By contrast, “steeper” regions of the landscape have higher ratchet robustness, since selection coefficients are larger. These considerations seem to suggest an intrinsic tension between mutational robustness and ratchet robustness. Ratchet robustness can also be realized by increased ratio of beneficial to deleterious mutation rates, represented on the fitness landscape by the increased probability of going “uphill” rather than “downhill” when locally exploring the genotype space via mutations (Goyal et al. 2012).

The goal of this paper is to identify those features of the fitness landscape that are favored by natural selection as a consequence of the fact that most mutations are deleterious. We investigate, in isolation, the effect of selection coefficient, and the influence of ratio of beneficial to deleterious mutation rates. We find that ratchet robustness protects populations from extinction due to deleterious mutations and should therefore be favored in the long term, although in the short term mutational robustness can also be selected for.

## Methods

### Evolutionary model

For all simulations, we implement discrete-time Wright-Fisher evolutionary model with custom Python code. Within populations of fixed size *N*, each individual’s genotype is solely identified by the number of deleterious mutations it has, which is denoted by *i, i* = 0,1…, *L. L* represents the genome length, which is also the maximum number of deleterious mutations possible. Populations are represented by vectors *V* of length *L* + 1, where each element *V_i_* records the number of individuals with *i* deleterious mutations so that ∑_*i*_ *V_i_* = *N*. Individuals with *i* mutations have fitness denoted by

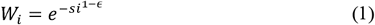

with selection coefficient s and epistasis parameter *ϵ. ϵ* = 0 in the absence of epistasis (**Fig. 1**), *ϵ* < 0 with negative epistasis (**Fig. 2A & B**: *ϵ* = −0.25), and *ϵ* > 0 with positive epistasis (**Fig. 2C & D**: *ϵ* = 0.25). Fitness defined by **Eqn. 1** depends only on the number of deleterious mutations in the genome and not their identity. We designate such fitness landscapes as isotropic.

**Figure 1:**
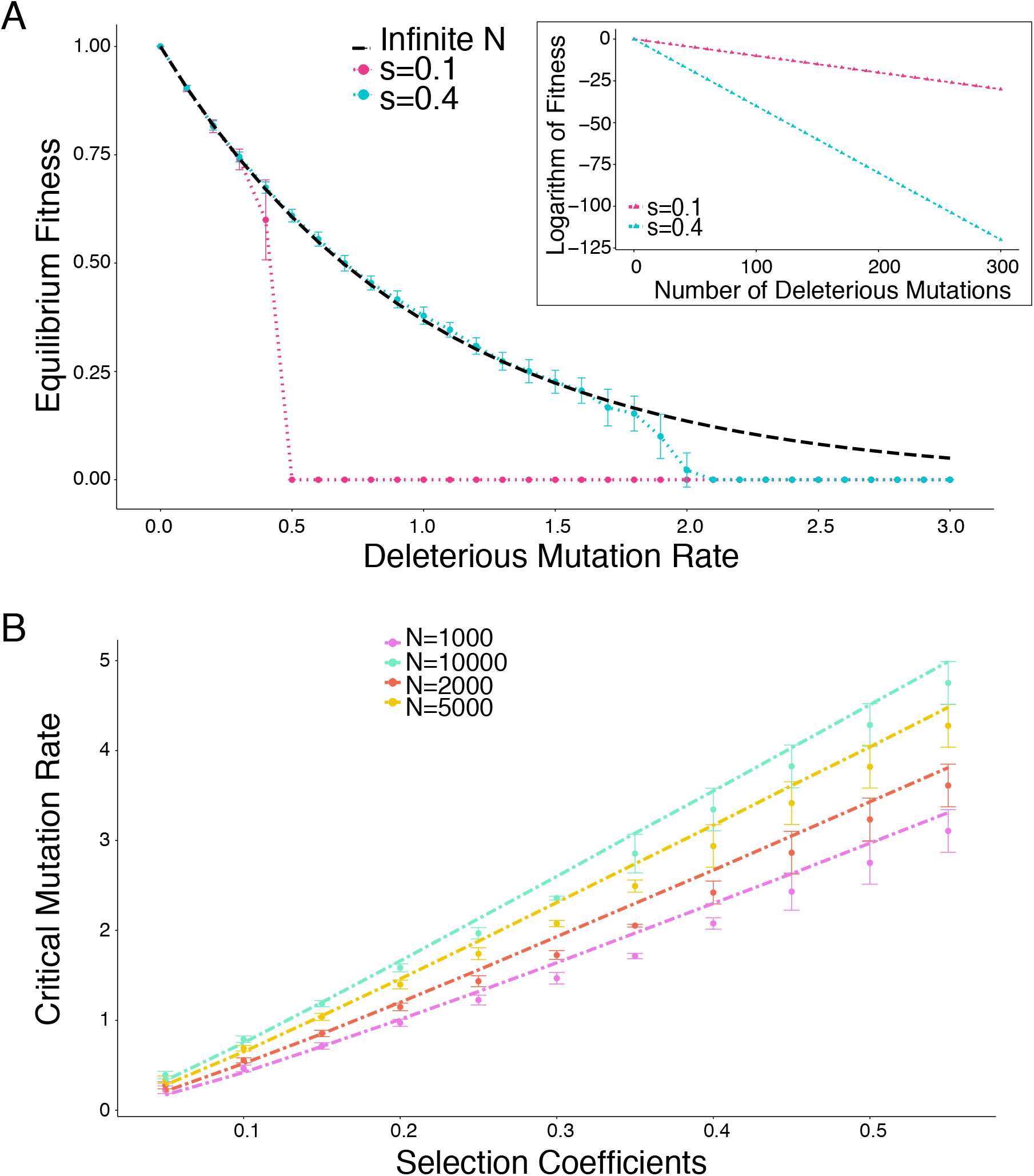
Under a fixed ratio of beneficial to deleterious mutation rates, populations residing on steeper fitness landscapes survive under higher deleterious mutation rate. A. Equilibrium fitness of finite populations under different deleterious mutation rates (*U_del_*) on fitness landscapes with two different selection coefficients (*s*). Dashed black line: theoretical prediction for equilibrium fitness in infinite populations (*e^−U_del_^*). Populations on the steeper fitness landscape (blue), i.e., the one with a larger s, maintain MSDE under higher *U_del_* than do populations on the shallower landscape (red). Beneficial mutation rate *U_ben_* = 0.01*U_del_*. Population size *N* = 1000. Average fitness is recorded after evolving for 10,000 generations (see **Methods**). Error bars: standard deviation over 50 replicates. Inset: comparison of the two fitness landscapes, each of which is composed of one isotropic peak without epistasis. B. Critical *U_del_* values from simulations (points) and analysis (lines, derived numerically from **Eqn. 7** in Goyal et al. 2012), for landscapes with different selection coefficients in the absence of epistasis. Error bars: uncertainty associated with locating the critical *U_del_* from simulation results (see **Methods**). Regardless of population size, populations on steeper landscapes always survive under higher *U_del_*.

Each generation of evolution consists of selection and mutation. The number of offspring with *i* mutations is a random variable proportional to *V_i_*, i.e., number of parents carrying *i* mutations, times *W_i_*, i.e., fitness of these parents. Then, mutations are imposed by sampling from Poisson distribution with parameter *U_dei_* to generate number of deleterious mutations, and from Poisson distribution with parameter *U_ben_* to generate number of beneficial mutations. Finally, a new vector *V*′ is constructed, where 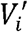 records number of individuals with *i* mutations in the new generation. Simulations are performed for a prespecified number of generations, usually 10,000, to ensure the result is unaffected by transient effects. We hold population size constant in all our simulations, i.e., we assume soft selection, and characterize populations in which all individuals are carrying *i* = *L* deleterious alleles as being at extinction.

### Infinite genome

For the first part of the results, we hold constant the ratio of beneficial to deleterious mutation rates (*U_ben_/U_del_* = 0.01). Conceptually this is equivalent to the well-known infinite sites model (Kimura 1969).

Simulations in **Fig. 1** and **Fig. 2** are performed on simple fitness landscapes with *L* = 300, with only one isotropic peak defined by **Eqn. 1**.

**Figure 2:**
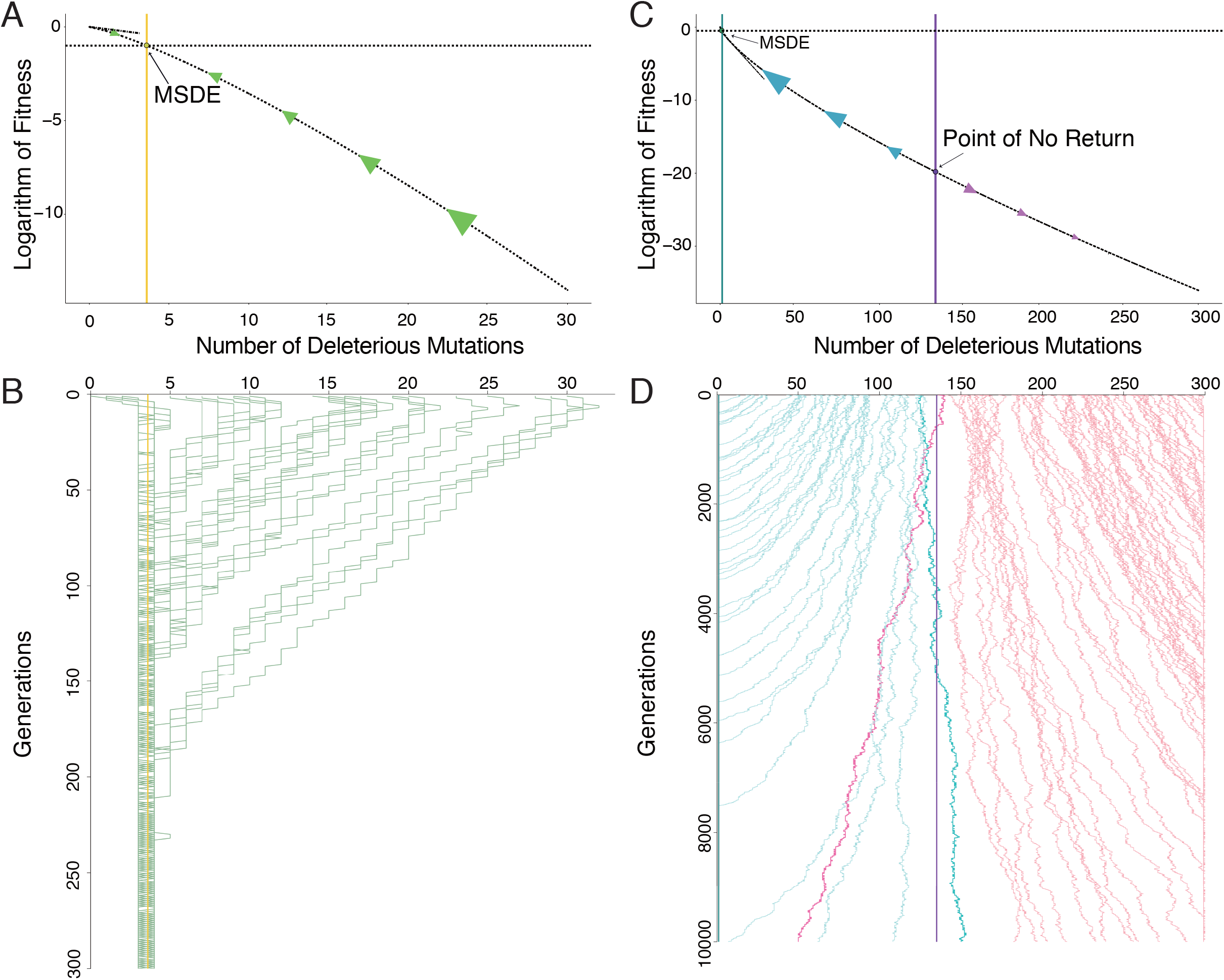
Fitness landscapes with negative epistasis protect populations from Muller’s ratchet while ones with positive epistasis make populations vulnerable to the ratchet. A. Cartoon of evolutionary dynamics of populations on fitness landscape composed of one isotropic peak with negative epistasis. Curved dashed line: fitness landscape. Straight tangent dashed line: fitness landscape without epistasis that shares the same selection coefficient at the peak. Note that the selection coefficient (the slope of the fitness landscape) increases with the number of deleterious mutations. Golden vertical line and horizontal dashed line: stable MSDE at the peak *e^−U_del_^* (see main text, also S1 Text). B. Time course of average number of deleterious mutations in 35 populations initialized at random points on isotropic fitness peak with negative epistasis during 10,000 generations (*N* = 1000, *s* = 0.2, *U_del_* = 1.0, *U_ben_* = 0.01*U_del_*, *ε* = −0.25). Golden vertical line: same as in A. All populations converge to the MSDE regardless of where they are initiated (green traces). C. Cartoon of evolutionary dynamics of populations on fitness landscape composed of one isotropic peak with positive epistasis. Curved dashed line: fitness landscape. Straight tangent dashed line: fitness landscape without epistasis that shares the same selection coefficient at the peak. Note that selection coefficient (equal to the slope of the fitness landscape) decreases with the number of deleterious mutations. Cyan vertical line and horizontal dashed line: unstable MSDE at the peak *e^−U_del_^* (see main text). Violet vertical line: predicted point of no return (derived numerically from **Eqn.7** in Goyal et al. 2012). D. Time course of average number of deleterious mutations in 100 populations initialized at random points on an isotropic fitness peak with positive epistasis during 10,000 generations (*N* = 1000, *s* = 0.5, *U_del_* = 0.5, *U_ben_* = 0.01*U_del_*, *ε* = −0.25). Cyan and violet vertical lines: same as in C. Populations initiated above the point of no return tend to evolve to the peak (blue traces), whereas ones initialized below it tend to succumb to Muller’s ratchet (pink traces). Realizations that fluctuate across the point of no return are colored in brighter blue and pink.

We characterize the critical *U_del_*, i.e., the highest *U_del_* under which populations can resist Muller’s ratchet with reference to the time variance in fitness. In **Fig. S2**, we present the variance in fitness across time at equilibrium for the simulations conducted in **Fig. 1A**. Notably, the critical *U_del_* i.e., the highest *U_del_* under which populations can resist Muller’s ratchet is seen to overlap with the *U_del_* under which time variance in fitness reaches maximum. This is reasonable, because when *U_del_* is well below or well above the critical *U_del_*, populations are either at MSDE or extinction and thus exhibit low time variance in fitness. In contrast, near the critical *U_del_*, the “tug-of-war” between the two possible states maximizes stochasticity.

We utilize the above observation to approximately locate the critical *U_del_* in **Fig. 1B**, across different population size *N* and selection coefficient *s*. For each *N* and *s* combination, we record the time variance in fitness under different *U_del_*, and report the *U_del_* associated with highest time variance as the critical *U_del_*. There are three sources of uncertainty in this analysis. First, it is impossible to conduct simulations under every possible *U_del_* value. In practice, we sample *U_del_* at intervals of 0.025, meaning that the “true” critical *U_del_* may fall in between two adjacent sampled *U_del_* values. Second, because of this intrinsic granularity, the detected maximum time variance may not be the true maximum. We measure such uncertainty as the range of *U_del_* values that show time variance above half of the detected maximum. Third, there is inevitable variation in recorded fitness variance across replicated simulations, due to the stochastic nature of the simulation. However, we find that the second source of variation dominates the other two by at least one order of magnitude (not shown). Therefore, we portray only the second type of uncertainty as error bars in **Fig. 1B**.

For simulations in **Fig. 3**, as shown in **Fig. 3A**, we fuse two isotropic peaks together: on the positive epistasis side of the intervening fitness valley, *ϵ* = 0.25 and *L* = 270; on the negative epistasis side, *ϵ* = −0.25 and *L* = 29. (These values of *L* were chosen so that fitness values on both sides of the floor of the valley would match.) Mutation on both sides of the valley were biased as described above, i.e. *U_ben_/U_del_* = 0.01.

**Figure 3:**
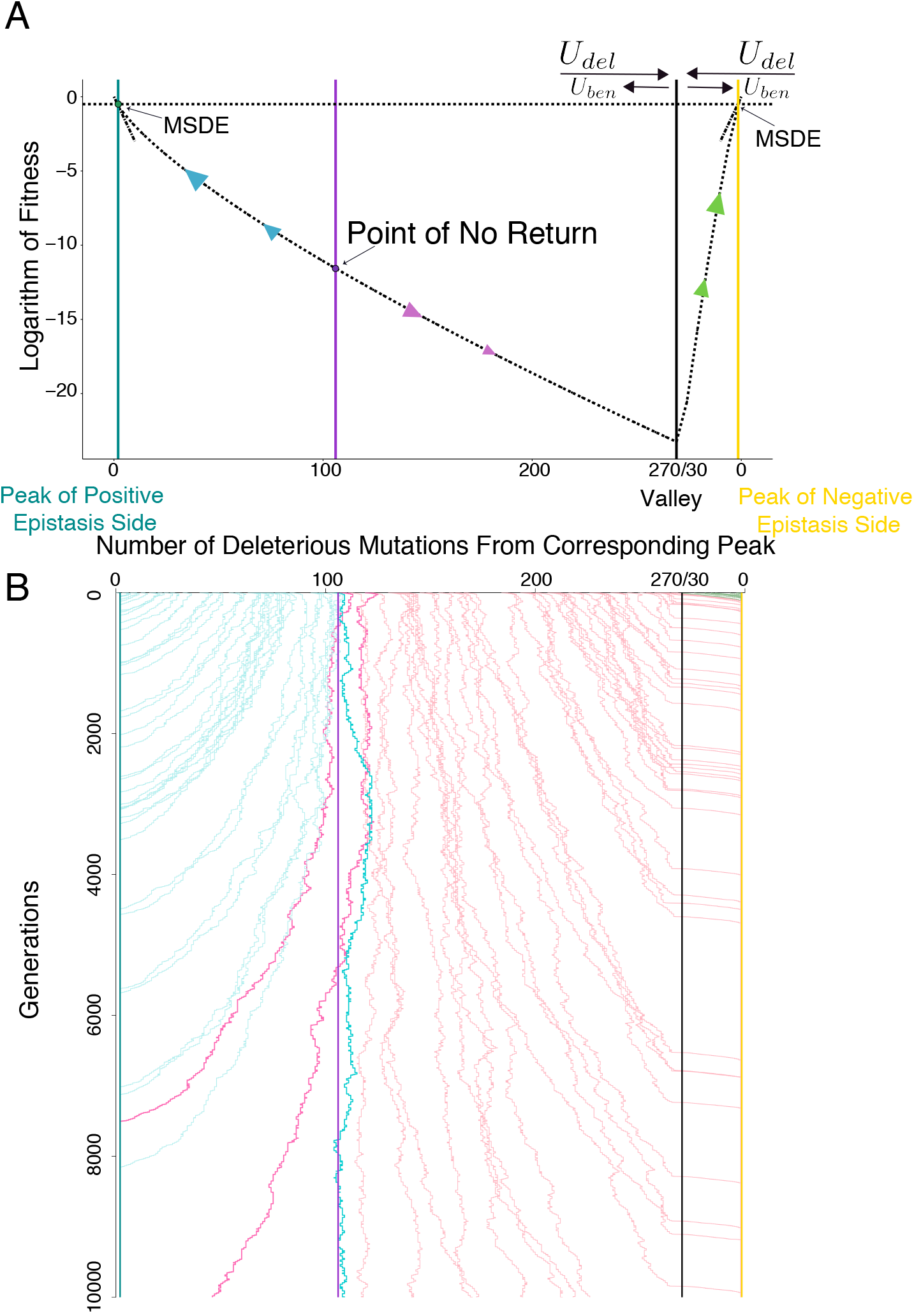
Populations converge to peak with negative epistasis on multi-peak fitness landscap. A. Cartoon of evolutionary dynamics of populations on multi-peak fitness landscape. Both fitness peaks are isotropic, one with only positive epistasis (left) and the other with only negative epistasis (right). Curved dashed lines on both sides represent the fitness landscape. Straight tangent dashed line on both sides again represent the slope (i.e. the selection coefficient at the peaks); this quantity is equal at both peaks. Horizontal dashed line: on both sides. Cyan vertical line: *e^−U_del_^* on the positive epistasis side. Violet vertical line: numerically derived point of no return on the positive epistasis side (**Eqn.7** in Goyal et al. 2012). Black vertical line: valley of the landscape. Golden vertical line: *e^−U_del_^* on the negative epistasis side. B. Time course of average number of deleterious mutations in 100 populations initiated at random, uniformly distributed points on multi-peak landscape over the course of 10,000 generations (*N* = 10,000, *s* = 0.3, *U_del_* = 0.5, *U_ben_* = 0.01 *U_del_*, positive epistasis side *ε* = 0.25, negative epistasis side *ε* = −0.25). Cyan, violet, black and golden vertical lines: same as in B. Populations are color coded based on their starting point (blue traces: above the point of no return on the positive epistasis side, pink traces: below the point of no return on the positive epistasis side, bright pink and blue traces: realizations that fluctuate across the point of no return on the positive epistasis side, green traces: populations initialized on the negative epistasis side). The vast majority of populations below the point of no return rapidly cross the valley and converge to MSDE on the negative epistasis side. Due to uniformly stronger selection on the negative epistasis side of the valley, populations there exhibit less stochasticity in their trajectories. Simulations in smaller population size (*N* = 1000) show qualitatively the same results, although it takes much longer to converge to the negative epistasis side (data not shown).

For simulations in **Fig. 4**, we define a hybrid fitness landscape consisting of two domains, one exhibiting positive epistasis and the other exhibiting negative epistasis. As shown in **Fig. 4A**, a fraction *p* of the first mutations put individuals in the positive epistasis domain, where *ϵ* = 0.25 and *L* = 300, while the remaining fraction 1 – *p* of first mutations put individuals at the negative epistasis domain where *ϵ* = −0.25 and L = 300. If a lineage returns to the peak (by beneficial mutations), the same *p* and 1 – *p* apply, i.e., lineages can “travel” between the two domains through the peak.

**Figure 4:**
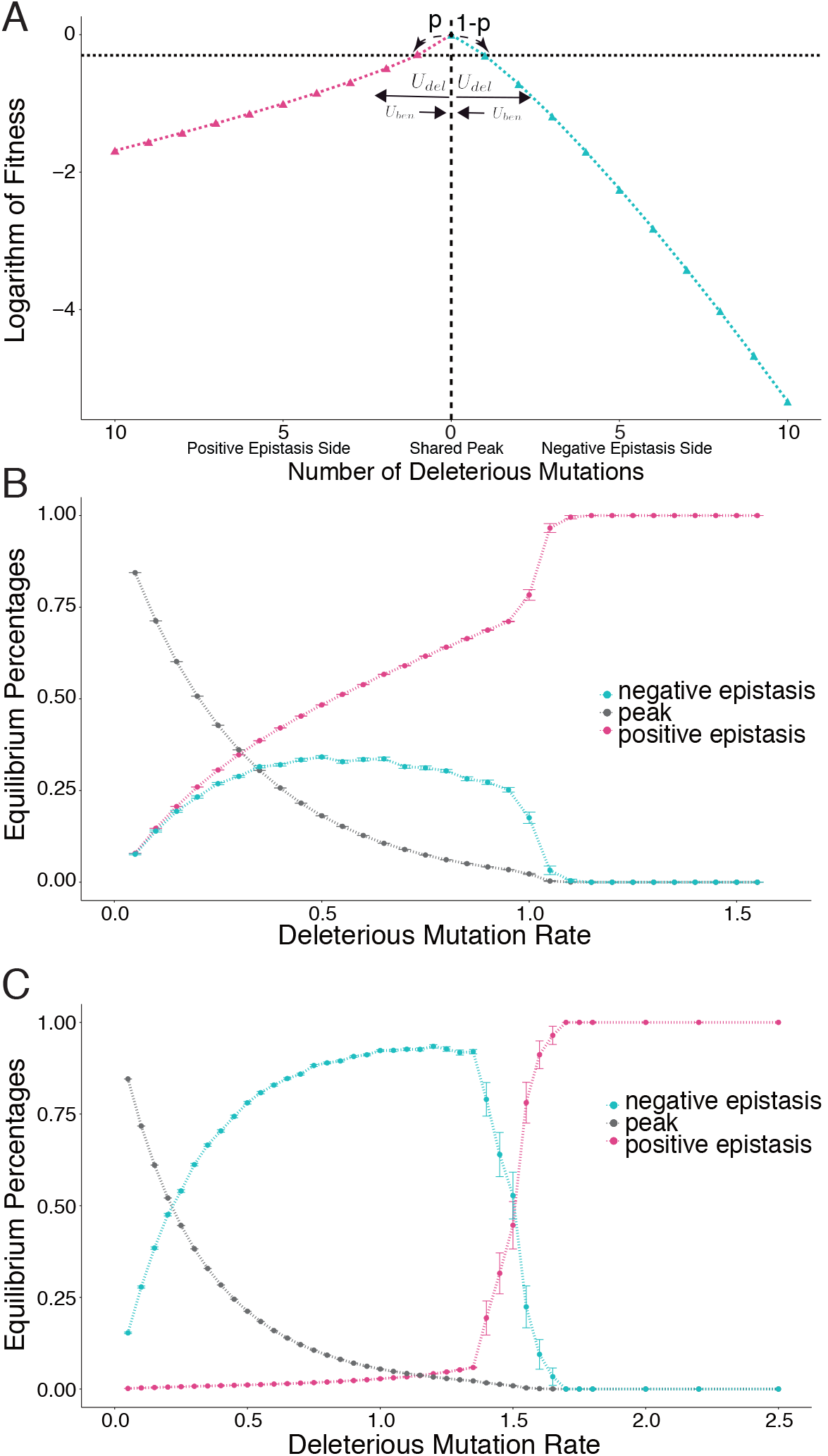
Mutational robustness and ratchet robustness on hybrid peaks. A. Cross-section through a hybrid fitness peak. While all first mutations in the highest fitness genotype share the same fitness effect, a fraction *p* of them cause subsequent mutations to exhibit positive pairwise epistasis, and the remaining fraction 1 – *p* cause subsequent mutations to exhibit negative epistasis. Populations are always initiated at the peak. Here *p* = 0.5. B. Equilibrium proportions of individuals at the peak (black), in the negative epistasis region (blue), and in the positive epistasis region (pink), as a function of *U_del_*(*U_ben_* = 0.01*U_del_, p* = 0.5, *N* = 1000, *s* = 0.35, negative epistasis region: *ε* = −0.25, positive epistasis region: *ε* = 0.25), evolved for 10,000 generations, error bars: standard deviation across 50 replicates. Under *U_del_* less than the critical *U_del_* at the peak (here ~1.1; see main text), the subpopulation on the negative epistasis region relies on continual mutational input from subpopulation on the peak. After *U_del_* exceeds the critical *U_del_* at the peak, the subpopulation on the peak goes extinct, and with it, the subpopulation on the negative epistasis region. At this point, the remaining population finds itself beyond the point of no return on the positive epistasis region (see main text), and it succumbs to Muller’s ratchet. Equilibrium proportions of individuals at the peak (black), the negative epistasis region (blue), and the positive epistasis region (pink), as a function of *U_del_* (parameters are exactly the same as panel B except *p* = 0.01), evolved for 10,000 generations, error bars: standard deviation across 50 replicates. Population dynamics resemble *p* = 0.5 (see main text).

### Finite genome

For the second part of the results, we allow *U_ben_/U_del_* to evolve with the proportion of deleterious mutation in the genome. Specifically, *U_ben_* equals proportion of deleterious mutations times the total mutation rate *U*, while *U_del_* equals *U* – *U_ben_*. Consequently, *U_ben_* and *U_del_* are unique for individuals in different *V_i_*, and we can only specify total mutation rate. In **Fig. 5**, simulations are conducted on an isotropic non-epistatic fitness landscape with *L* = 10 and *s* = 0.4.

**Figure 5:**
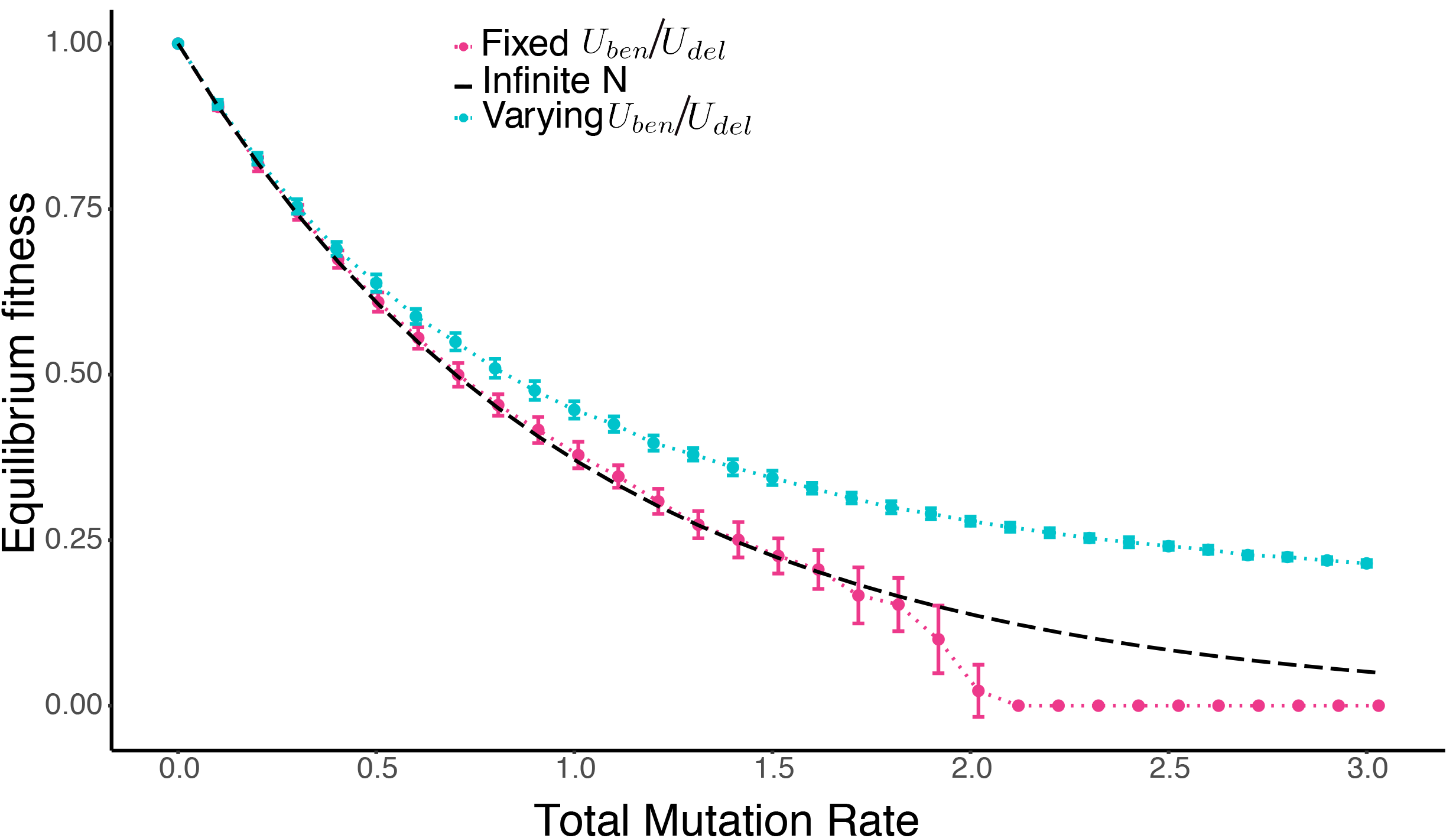
Increasing *U_ben_/U_del_* with decreasing fitness protects populations from Muller’s ratchet. Equilibrium fitness as a function of total mutation rates (*U*) in finite asexual populations with evolving and fixed *U_ben_/U_del_* Dashed black line: theoretical prediction for equilibrium fitness in infinite populations (*e^−U_del_^*) with fixed *U_ben_/U_del_*(*U_ben_* = 0.01*U_del_*). Population size *N* = 1000. Selection coefficient *s* = 0.4. Average fitness is recorded after evolving for 10,000 generations (see **Methods**). Populations in which *U_ben_/U_del_* evolve by virtue of reversion mutations (blue) survive under high *U*, while ones with fixed *U_ben_/U_del_*(red) succumb to Muller’s ratchet. (These latter results are identical to those shown for *s* = 0.4 in **Fig. 1**.) Error bars: standard deviation over 50 replicates.

## Results

We use individual-based simulations of finite populations of asexual organisms to study the ability of selection and evolving ratios of beneficial to deleterious mutation rates to protect populations from deleterious mutations. We first assume a constant ratio of beneficial to deleterious mutation rates and examine evolutionary behavior as a function of fitness landscape topography. We then allow the ratio of beneficial to deleterious mutation rates itself to evolve while holding other aspects of the fitness landscape constant.

### Effect of fitness landscape topology

We simulate evolution on four kinds of fitness landscapes with increasing complexity, while holding the ratio of beneficial to deleterious mutation (*U_ben_/U_del_*) equal to 0.01. We define a fitness peak as isotropic if the fitness of any genotype only depends on its number of deleterious mutations, but not the identity of these mutations. We start by examining evolutionary behavior on fitness landscapes with a single peak where every deleterious mutation shares the same effect regardless of the current genome (**Eqn. 1**: *ϵ* = 0, **Fig. 1A** inset). This is, by definition, an isotropic peak in the absence of epistasis. Next, we allow the effect of deleterious mutations to increase (**Eqn. 1**: *ϵ* < 0, **Fig. 2A**) or decrease (**Eqn. 1**: *ϵ* > 0, **Fig. 2C**) with the number of deleterious mutations in the current genome. In other words, we are adding epistasis to the isotropic peak, distinguishing between negative (**Fig. 2A**) and positive (**Fig. 2C**) epistasis. We then examine evolutionary behavior on fitness landscapes consisting of mutationally adjacent isotropic peaks with negative and positive epistasis (**Fig. 3A**). Finally, we relax our assumption of isotropic peaks, and model populations evolving on hybrid fitness peaks characterized by domains of positive and negative epistasis (**Fig. 4A**).

#### In the absence of epistasis, populations residing on steeper fitness landscapes survive under higher deleterious mutation rate

We first confirm the effect that stronger selection is better able to protect populations from Muller’s ratchet, or equivalently, that mutational robustness can enable the ratchet (Goyal et al. 2012). To do so, we construct two fitness landscapes, each composed of one isotropic peak but with different steepness (selection coefficients *s*) in the absence of epistasis (**Fig. 1A** inset). We simulate the evolution of finite asexual populations on such landscapes under different deleterious mutation rates, and record their fitness at equilibrium (**Fig. 1A**). As mentioned above, we implement soft selection, and we define extinction as occurring when all individuals in the population attain the minimum possible fitness.

Under low mutation rates, selection is able to purify deleterious mutations, and drift (and thus, the ratchet) is irrelevant on both flat and steep landscapes. This is because the stochastic loss of fittest class is extremely rare: purifying selection is strong relative to mutation, and if lost, the fittest class is quickly restored by beneficial mutations. Consequently, equilibrium fitness lies close to the well-known infinite population expectation *e^−U_del_^* (black dashed line in **Fig. 1A**, Haldane 1937; Kimura and Maruyama 1966). However, after *U_del_* exceeds some critical value (at this population size, *U_del_* ≈ 0.4 for *s* = 0.1 and *U_del_* ≈ 1.9 for *s* = 0.4, see **Fig. 1A**), rapidly accumulating deleterious mutations overwhelm selection, leading to extinction by Muller’s ratchet. We name the highest *U_del_* under which populations are able to resist Muller’s ratchet the “critical *U_del_*”. Our simulation results for the critical *U_del_* under different *N* and *s* are consistent with analytical results based on **Eqn. 7** in Goyal et al. 2012 (see **Fig. 1B**). Given a fixed population size (*N*) and constant *U_ben_/U_del_*, populations on steeper landscapes are able to survive under higher *U_del_*, i.e., they demonstrate a higher critical *U_del_* (**Fig. 1A & B**, Goyal et al. 2012). In other words, in the long term, increased mutational robustness increases susceptibility to Muller’s ratchet.

#### Fitness landscapes with negative epistasis protect populations from Muller’s ratchet while ones with positive epistasis make populations more vulnerable to the ratchet

Epistasis means the dependence of mutational fitness effects on the genetic background. Without epistasis (**Fig. 1**), there is no evolution of selection coefficients, since each mutation will always have the same effect. Put differently, epistasis determines how selection changes as deleterious mutations accumulate, shaping both local mutational and ratchet robustness experienced by an evolving population. In order to understand the role of epistasis, we consider the two simplest cases: one where selection strength monotonically increases with the number of deleterious mutations, and the other where selection strength monotonically decreases with the number of deleterious mutations. In other words, we consider a single isotropic peak with either only negative epistasis (**Fig. 2A**) or only positive epistasis (**Fig. 2C**), respectively.

When epistasis is negative, as deleterious mutations accumulate, the local strength of purifying selection increases, and consequently subsequent deleterious mutations become less likely to accumulate. This hints at a negative feedback between the accumulation of deleterious mutations and the tendency to accumulate more, which potentially could halt Muller’s rachet (as previously seen in Kondrashov 1994). On the other hand, when epistasis is positive, as deleterious mutations accumulate, local selection weakens, and deleterious mutations can more easily accumulate. This suggests a positive feedback between the aforementioned two forces, which may render populations more vulnerable to the ratchet than those examined above. Importantly, the critical *U_del_* will thus vary across the fitness landscape in response to this variation in the strength of selection

Our simulations are consistent with this intuition (**Fig. 2B, Fig. 2D**). In the presence of negative epistasis, when *U_del_* is lower than the critical *U_del_* at the peak, it is lower than the critical *U_del_* anywhere on the landscape. Therefore, selection will drive the population back toward the peak regardless of where on the landscape it is initialized (**Fig. 2B**). Even if *U_del_* is higher than the critical *U_del_* at the peak, there exists a point on the landscape where selection exactly offsets such *U_del_*, because purifying selection increases monotonically from the peak (see also **S1 Text**). If a population is initialized below this point, selection locally will be strong enough to push populations upward until this point is reached. If instead, a population is initialized above this point, mutation will be strong enough to push the population downward to this point, but no further. Therefore, even under high *U_del_*, landscapes with negative epistasis will halt Muller’s ratchet and establish stable MSDE.

Conversely, on the landscape with positive epistasis, when is lower than the critical at the peak, populations initiated at the peak can maintain MSDE. However, positive epistasis means that purifying selection decreases monotonically from the peak, and consequently there exists a point on the landscape where selection exactly offsets such. Populations initiated above this point will experience selection stronger than required to offset and evolve to the peak, while populations initiated below this point suffer from selection weaker than needed to offset *U_del_*, and will succumb to Muller’s ratchet (**Fig. 2D**). Therefore, we refer to this point as “point of no return”. Importantly, even if parameter values are such that populations can equilibrate at the peak, stochastic fluctuations will eventually take them across this point of no return, after which they will succumb to the ratchet. Moreover, as *U_del_* increases, the selection required to oppose mutation naturally increases as well. Correspondingly, as increases, the point of no return migrates towards the peak. This imposes a greater danger of succumbing to Muller’s ratchet for populations in the vicinity of the peak via stochastic fluctuations. Finally, once *U_del_* is greater than the critical *U_dei_* at the peak, the point of no return overlaps the peak, and mutation overwhelms selection everywhere on the fitness landscape.

#### Populations converge to the peak with negative epistasis on multi-peak fitness landscape

We next demonstrate that populations can indeed evolve to occupy regions of the fitness landscape with negative epistasis when challenged by deleterious mutations. To do so, we construct a fitness landscape composed of two mutationally adjacent isotropic peaks featuring opposite signs of epistasis (**Fig. 3A**). Populations finding themselves below the point of no return on the positive epistasis side will experience selection weaker than needed to offset mutation and their fitness will decline to the valley, similar to populations declining to the bottom of the landscape in **Fig. 2D**. Moreover, populations above the point of no return will nevertheless experience stochastic fluctuations and will be eventually carried over the point of no return, at which point they will also evolve to the valley. However, upon arrival at the valley, strongly beneficial mutations become available, drawing populations onto the negative epistasis side of the valley, after which they quickly climb to MSDE (**Fig. 3B**). (Note that *U_ben_* = 0.01 *U_del_* on both sides of the valley, and consequently this behavior is driven entirely by differences in the local strength of natural selection.) Since the two peaks share identical selection coefficients at the peak, the positive epistasis side has uniformly higher mutational robustness but uniformly lower ratchet robustness. This extends our understanding of the tension between mutational and ratchet robustness and demonstrates that ratchet robustness, instead of mutational robustness, is likely to evolve in response to high deleterious mutation rates.

#### Mutational robustness and ratchet robustness on hybrid peaks

Biologically realistic landscape peaks are unlikely to have only negative or positive epistasis. As a first attempt to investigate the complexity introduced by heterogeneity in epistasis, we now allow the sign of epistatic effects of mutations to be dependent on the first mutation away from the peak. Specifically, a certain fraction (*p*) of the first mutations now place the evolving population on a region of the landscape exhibiting positive epistasis, while the remaining fraction (1 – *p*) place the evolving population on a region of the landscape exhibiting negative epistasis (**Fig. 4A**). Concretely, among all the mutational paths leaving the peak, a proportion *p* of them show positive epistasis, while the remaining fraction 1 – *p* show negative epistasis. We assume that all first mutations share identical fitness effects, so that there is no immediate fitness advantage to lineages entering the region with negative or positive epistasis. However, for all genotypes with greater than one deleterious mutation, fitness is necessarily higher in regions of the landscape with positive epistasis than in regions with negative epistasis (**Eqn. 1**). In other words, regions of the landscape with positive epistasis have both higher fitness and higher mutational robustness (but lower ratchet robustness) compared with ones with negative epistasis. Note that the only mutational path between regions is through the peak. We evolved populations on such hybrid peak with *p* = 0.5 and report the proportions of populations residing exactly at the peak, in the negative epistasis region, and in the positive epistasis region as a function of *U_del_* (**Fig. 4B**).

When *U_del_* is so low that majority of the individuals carry zero or one mutation (here *U_del_* ≈ 0.2, **Fig. 4B, Fig. S3A**), the proportion of the population on the negative epistasis region is very close to that on the positive epistasis region. This merely reflects the fact that *p* equals 0.5, since the fitness cost of the first mutation is the same. However, as *U_del_* rises to moderate level (here ~0.2 < *U_del_* < ~1.1, **Fig. 4B, Fig. S3B**), the proportion of the population on the negative epistasis region begins to drop below 0.5. This reflects the selective enrichment of the subpopulation experiencing positive epistasis: individuals with two or more mutations on the positive epistasis region are fitter than are those with the same number of mutations on the negative epistasis region. Nevertheless, in this intermediate range of values of *U_del_*, a subpopulation on the negative epistasis region is still sustained despite lower fitness, thanks to net mutational inflow from the subpopulation at the peak (**Table S1**). In essence, the subpopulation on the negative epistasis region is at mutation-selection balance: constantly being purified by selection but being regenerated by mutation from the peak.

However, after *U_del_* increases enough that the peak can no longer be sustained (here, *U_del_* ≈ 1.1), the proportion of the population at the peak become negligible (**Fig. 4B, Fig. S3C**). As a result, the subpopulation residing on the negative epistasis region of the fitness landscape is mutationally disconnected from the peak and is quickly wiped out by selection. The remaining population now occupies only the positive epistasis region. However, since *U_del_* has overwhelmed selection at the peak, it necessarily does so also at every other point in the positive epistasis region, and the population quickly succumbs to Muller’s ratchet. This threshold recapitulates results seen when the point of no return reaches the peak on isotropic fitness peaks with positive epistasis region (**Fig. 2C & 2D**, although the numeric value of the critical mutation rate differs here, reflecting its dependence on the fitness landscape).

More importantly, the observed pattern does not depend on the particular value *p* = 0.5: even when there is only a very small fraction of paths leaving the peak with positive epistasis, subpopulations on the positive epistasis region of the landscape will be favored due to their short-term fitness advantage (**Fig. 4C**). Such an advantage is amplified by higher *U_del_*, so long as it remains less than the critical *U_del_* at the peak, i.e., the *U_del_* under which individuals at the peak are lost to Muller’s ratchet. Below this threshold, the two subpopulations accumulate more mutations, and thus experience increased fitness differences. Eventually, once *U_del_* exceeds the critical *U_del_* at the peak, the mutational connection between subpopulations on the fitness landscape is again extinguished. At this point both subpopulations are doomed. The lower-fitness subpopulation on the region of the landscape with negative epistasis will lose its mutational input and go extinct. At the same time, the remaining higher-fitness subpopulation on the region with positive epistasis will succumb to Muller’s ratchet.

### Effect of ratio of beneficial to deleterious mutation rates

In order to isolate the role that topographic features of the fitness landscape play in an evolving population’s response to deleterious mutations, we have thus far assumed that the ratio of beneficial to deleterious mutation rates remains fixed (see **Methods**). However, as deleterious mutations accumulate, this ratio can itself evolve, due to reversion mutations. Moreover, compensatory mutations (mutations that are only beneficial in the presence of deleterious mutations at other loci) amplify this effect. For example, one survey estimated that each deleterious mutation gives rise to approximately 12 compensatory mutations (Poon and Chao 2005).

#### Increasing ratio of beneficial to deleterious mutation rates with decreasing fitness protects populations from Muller’s ratchet

We therefore modify our simulations to allow deleterious mutations to revert, thereby causing *U_ben_/U_del_* to increase with the accumulation of deleterious mutations. Biologically, this approach is conservative because it neglects compensatory mutations induced by deleterious mutation (e.g., Poon and Chao 2005). Since increasing *U_ben_/U_del_* reduces the rate at which new deleterious mutations occur, this suggests that another negative feedback may exist between the accumulation of deleterious mutations and the tendency to accumulate more (Goyal et al. 2012). This is reminiscent of the one due to negative epistasis seen above (**Fig. 2A**). To gain more insight into this effect, we compare equilibrium fitness in populations evolved when is allowed to change with that seen in populations evolved under fixed *U_ben_/U_del_* (**Fig. 5**). As *U* increases and deleterious mutations accumulate, we see that populations with varying enjoy the benefit of lower *U_del_* and equilibrate at higher fitness than do populations with fixed *U_ben_/U_del_*. Eventually, under sufficiently high *U*, populations with fixed *U_ben_/U_del_* succumb to Muller’s ratchet, as seen above. In contrast, populations with varying *U_ben_/U_del_* halts the ratchet, as previously described (Goyal et al. 2012).

## Discussion

### Negative feedbacks halt Muller’s ratchet

Our results highlight the importance of negative feedbacks in halting Muller’s ratchet. The mechanisms responsible for such negative feedback may be varied, realized via negative epistasis (**Fig. 2 & 3**), or increasing *U_ben_/U_del_* (**Fig. 5**). We note that many surveys of biological fitness landscapes find extensive evidence for negative epistasis (Bershtein et al. 2006; Costanzo et al. 2010; Khan et al. 2011; Steinberg and Ostermeier 2016). Additionally, *U_ben_/U_del_* tends to increase as deleterious mutations accumulate due to reversions and compensatory mutations. More importantly, both negative epistasis and increasing *U_ben_/U_del_* can be seen as representative cases of changing distribution of fitness effects (DFE) (Eyre-Walker and Keightley 2007) with decreasing fitness. Negative epistasis influences the magnitude of mutational effects, while increasing *U_ben_/U_del_* influences the ratio between beneficial and deleterious mutations. While outside the scope of our simulations, we suggest that as fitness goes down, both aspects of the DFE can simultaneously evolve, resulting in a wide range of potential DFE changes that confer ratchet robustness.

In fact, even in the presence of positive epistasis, it’s plausible that increases in *U_ben_/U_del_* could still halt Muller’s ratchet. Conversely, even if *U_ben_/U_del_* decreases as deleterious mutations accumulate, sufficiently strong negative epistasis could in principle protect populations from deleterious mutations. (Although as noted above, there is little reason to believe that *U_ben_/U_del_* should decline with fitness.) At this moment, we remain ignorant about the general rules driving changing DFE as a function of fitness. In the future, theoretical models that parameterize the DFE could be built to quantify the relative importance of negative epistasis and increasing *U_ben_/U_del_* in protecting evolving populations from deleterious mutation. Experimental studies could also be conducted to compare DFE at different fitness levels for various biological systems. Notably, a previous experimental study using bacteriophage has observed that as fitness declines, the rate of beneficial mutations changes much more drastically than does the size of mutational effects (Silander et al. 2007).

### Widespread existence of mutational robustness may result from selection for environmental robustness

Our results suggest that mutational robustness cannot be selected for the long term, and we address previous theoretical studies on the evolution of mutational robustness in **Box 2**. Nevertheless, mutational robustness is seen at many levels of biological systems (Wagner 2013). We note however that the existence of mutational robustness need not imply selection for mutational robustness (Hermisson and Wagner 2004; Proulx and Phillips 2005; Siegal and Leu 2014). Following others (de Visser et al. 2003), we suggest instead that mutational robustness may often evolve as a correlated consequence of selection for environmental robustness, i.e., an organism’s ability to sustain fitness against environmental perturbations. Theoretically and empirically, environmental robustness has been shown to give rise to mutational robustness (Wagner et al. 1997; Ancel and Fontana 2000; Meiklejohn and Hartl 2002; de Visser et al. 2003; Szollosi and Derenyi 2007; Goldsmith and Tawfik 2009; Domingo-Calap et al. 2010).

#### Box 2: Reconciliation with the literature on the evolution of mutational robustness

On close inspection, **Fig. 2** in Wilke et al. 2001 (reproduced here as **Fig. 6**) may suggest the evolution of ratchet robustness rather than of mutational robustness. Specifically, the fitness distribution of genotypes in the mutational neighborhood of the lineage adapted under high mutation rate appears to contain more lethal mutations than do those mutationally adjacent to the lineage adapted to low mutation rate. Furthermore, the lineage adapted under low mutation rate seems more prone to mutation accumulation than does the lineage adapted under high mutation rate. Both observations suggest that the lineage adapted under low mutation rate experiences weaker purifying selection than does the one adapted under high mutation rate. Thus, we interpret the data in Wilke et al. to suggest that the lineage adapted under high mutation rate actually has lower mutational robustness, and consequently higher ratchet robustness, consistent with our results.

Unfortunately, the two studies (Codoner et al. 2006; Sanjuan et al. 2007) attempting to experimentally validate conclusions in Wilke et al. failed to fully characterize the different lineages’ fitness landscapes. Specifically, only non-lethal genotypes were considered, while lethal mutations were ignored. Thus, we are unable to assess whether the lineage evolved under high mutation rate has higher mutational robustness than the lineage evolved under low mutation rate, since only a fraction of the DFE was measured.

Findings in van Nimwegen et al. (1999) are actually in agreement with our conclusions here, as previously noted (Wylie and Shakhnovich 2011). Specifically, by construction Muller’s ratchet is impossible in van Nimwegen et al.’s model, since any non-neutral mutation is lethal and so is instantaneously eliminated. Put another way, under this model evolving populations enjoy the benefits of increased mutational robustness without the otherwise concomitant risk of succumbing to Muller’s ratchet. Relaxing van Nimwegen et al.’s assumption of strict lethality for all non-neutral mutations recovers exactly our predicted behavior: at equilibrium, mutational robustness declines with mutation rate (see Fig S5 in Wylie and Shakhnovich 2011). The same argument also applies to other studies where all deleterious mutations are lethal, e.g., Wilke 2001.

**Figure 6:**
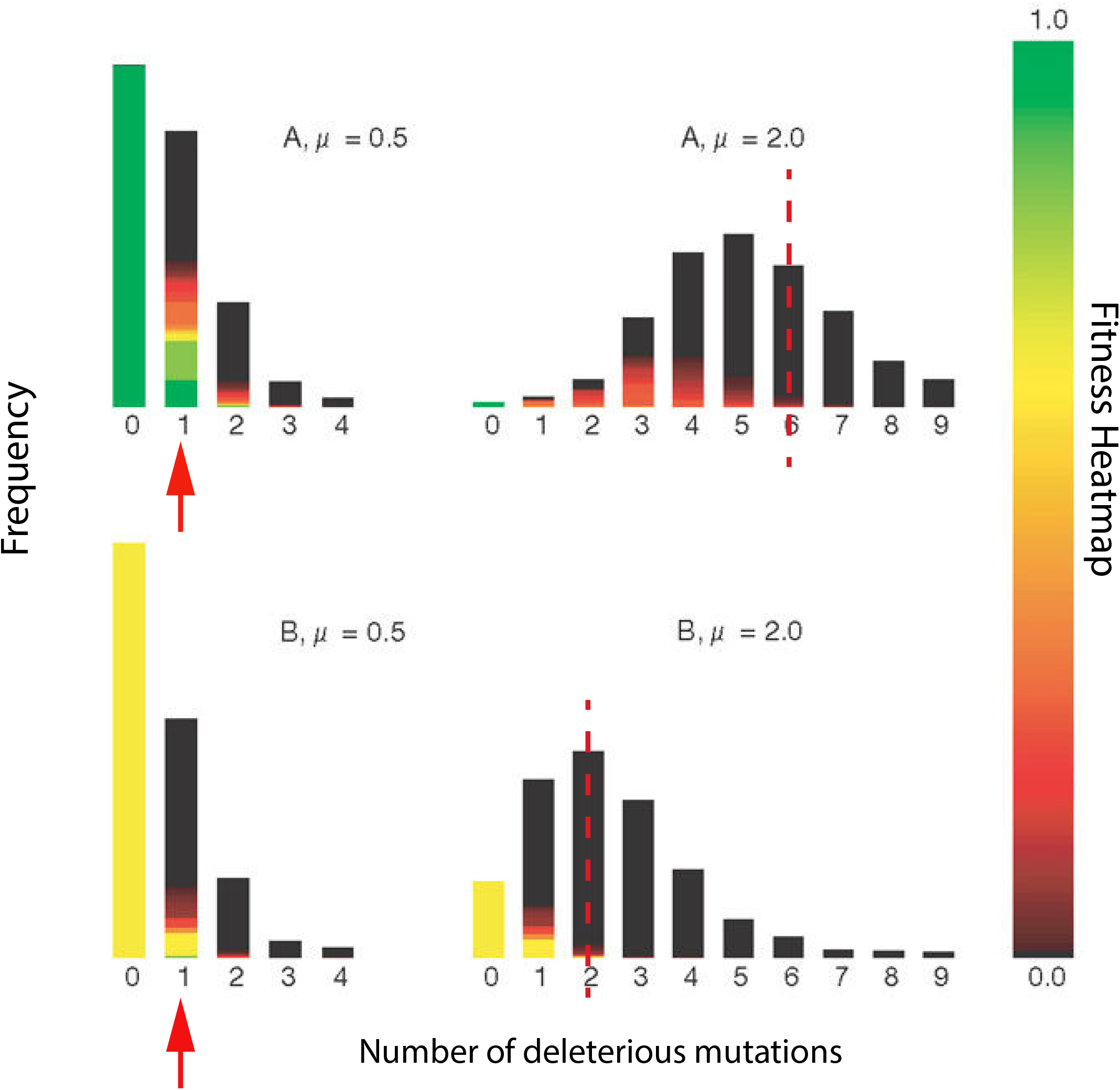
Annotated **Fig. 2** from Wilke et al. (2001) suggests the evolution of ratchet robustness in populations of digital organisms. Populations in Avida (Ofria et al. 2009) were evolved for 1,000 generations at low and high mutation rates (*μ* = 0.5 and 2.0, respectively). In this figure, the fitness landscapes in the mutational neighborhoods of these two populations were examined by propagating new populations seeded with one representative individual from each for 15 generations. Top panels: “A” designates new populations seeded with a representative individual adapted under low mutation rate. Bottom panels: “B” designates new populations seeded with a representative individual adapted under high mutation rate. *μ* denotes the mutation rate during the subsequent 15 generation experiment. Each bar indicates the frequency of genotypes with the corresponding number of mutations at the end of these 15 generations; colors represent the fitness of these genotypes. The left two panels illustrate the fitness landscape in the nearby mutational neighborhood of A or B (small *μ* = 0.5). Importantly, the descendants of the individual from the B population include many more lethal mutants (black) than do those from the individual from the A population (red arrows). The right two panels depict the fitness landscape across a broader mutational neighborhood of A or B (large *μ* = 2.0). Notably, the viable descendants of the individual from the B population carry fewer deleterious mutations than do those from the A population (red dashed lines represent the approximate mutational limit among viable offspring: upper panel skews towards more mutations than does the lower panel). Both observations suggest that selection against deleterious mutation is stronger in the mutational neighborhood occupied by the B population than it is in the neighborhood of the A population, consistent with the evolution of ratchet robustness rather than mutational robustness in response to elevated mutation rate.

Why can natural selection favor the evolution of environmental but not mutational robustness? The key distinction is that mutational robustness requires tolerating heritable perturbations, which inevitably alters the “starting point” of future generations. Such heritable decay is intrinsic to Muller’s ratchet. By contrast, selection for environmental robustness entails non-heritable environmental perturbation. Consequently, the short-term advantage of environmental robustness is not offset by any long-term cost, accounting for the absence of an “environmental ratchet”. In summary, while mutational robustness may be widespread in nature, we suggest one alternative interpretation for its evolution: namely as a correlated consequence of selection for environmental robustness (de Visser et al. 2003). We address another possibility next.

### Long-term and short-term fate of mutational and ratchet robustness

Our findings also illustrate that mutational robustness can be selected for in the short term, despite the danger it imposes in the long term (**Fig. 4**). This suggests another mechanism for the existence of mutational robustness in natural populations. How populations overcome this shortsightedness remains an open question. One possibility described in a previous study (O’Fallon et al. 2007) is that subdivision can protect populations from myopic selection for mutational robustness. Because selection is more effective at purging deleterious mutations in demes dominated by ratchet robust individuals, in that study net dispersal rates were higher from those demes, and the population in total was thus enriched for such individuals in spite of the short-term disadvantage. We predict that any population structure capable of hindering rapid fixation of mutational robustness will help natural selection favor ratchet robustness. However, a detailed survey of possible mechanisms is outside the scope of this study.

## Conclusion

It is of fundamental interest to understand how natural populations evolve to cope with the fact that almost all mutations are deleterious. Intuitively, natural selection may favor mutational robustness, as deleterious mutations of lesser effects seem to pose less danger. However, we have extended work of others to demonstrate another peril of deleterious mutations: long term extinction via Muller’s ratchet. Indeed, we show that mutational robustness increases the risk of Muller’s ratchet. We then demonstrate that both negative epistasis and increasing *U_ben_/U_del_* confer ratchet robustness and can protect populations from deleterious mutations in the long term. Overall, our findings indicate that natural selection can favor ratchet robustness over mutational robustness, and we define important areas of future theoretical and experimental work.

